# Autophagy induction by cytoplasmic Acinus requires Lrrk activity

**DOI:** 10.64898/2026.11.16.767156

**Authors:** Nilay Nandi, Charles Tracy, Helmut Krämer

## Abstract

Autophagy is a critical pathway for maintaining cellular health in ageing and disease but mechanisms matching levels of autophagy to different stress conditions remain incompletely understood. In response to proteostasis stress, autophagy is induced upon activation of Acinus, a mostly nuclear protein previously linked to alternative splicing as a subunit of the ASAP complex. Here, we show that Acinus induces autophagy independently of its function in splicing. Instead, autophagic flux is promoted by mutations of Acinus that either disrupt its interaction with the ASAP complex or otherwise increase its cytoplasmic fraction. Proximity labeling of the proteome preferentially associated with cytoplasmic rather than nuclear Acinus identified Rab32. Genetic analysis revealed that the Rab32 GTPase and its effector Lrrk kinase are required for autophagy induction by activated Acinus, in contrast to Lrrk-independent mTor-regulated autophagy. This specific requirement in Acinus-regulated autophagy has possible implications for Parkinson’s disease for which mutants in the human orthologs of Rab32 and Lrrk constitute risk factors.

## Introduction

Autophagy (here short for macroautophagy) is a conserved stress response pathway that can be activated by starvation (Ohsumi, 2014), damaged organelles or proteostasis stress with profound medical impact on conditions ranging from cancer and inflammation to neurodegeneration (Nixon and Rubinsztein, 2024; Stavoe and Holzbaur, 2019). Basic steps of the autophagy pathway include initial formation of phagophores, elongation of isolation membranes, their closure forming autophagosomes to fuse with late endosomes or lysosomes and deliver cargo for its degradation (Vargas et al., 2022). How these processes are modulated by different signaling pathways to match diverse physiological stress conditions in distinct cell types remains an important question (Wei et al., 2024).

One such regulatory pathway depends on the activation of Acinus (Acn) by the CDK5/p35 kinase complex (Nandi et al., 2017). Stressors that engage this pathway include aggregated polyQ proteins or Aβ peptides, and in turn, autophagy activated by Acn can reduce the levels of such aggregates and promote longevity (Nandi et al., 2014; Nandi et al., 2017). Here, we refer to such proteostasis stress-induced, but starvation and mTor inhibition independent autophagy as "basal autophagy". A role for Acn in regulating basal autophagy contributing to cellular quality control has emerged from genetic screens in *Drosophila* (Haberman et al., 2010) and mammalian cells (Orvedahl et al., 2011). The mechanism by which Acn elevates autophagy is however not well understood.

Acn and its mammalian Acin1 orthologs are mostly nuclear proteins that interact with the binding partners Sap18 and RNPS1 to form the conserved ASAP complex (Murachelli et al., 2012; Schwerk et al., 2003). Through its interactions with the Exon-Junction-Complex (Tange et al., 2005), the ASAP complex affects aspects of mRNA metabolism, including alternative pre-mRNA splicing in flies (Hayashi et al., 2014; Malone et al., 2014) and mammalian cells (Deka and Singh, 2019; Rodor et al., 2016; Wang et al., 2015).

Here, we address the mechanism for Acn-mediated regulation of autophagy. We find that the role of Acn in autophagy is independent of its function in splicing. Instead, autophagy activation by Acn depends on the cytoplasmic subset of Acn proteins. Proximity labeling of the proteome preferentially associated with cytoplasmic Acn identified Rab32 and its effector Lrrk kinase as major regulatory node. The requirement of Lrrk kinase in Acn-induced autophagy contrasts with it being dispensable for starvation-induced mTOR-regulated autophagy.

## Results

### Splicing function of Acn is not required for enhancing autophagy

Acn functional domains and regulatory elements are highly conserved (Fig. 1 A). We previously have generated transgenic flies in which endogenous Acn was replaced by either N-terminally Myc-tagged Acn^WT^, the caspase cleavage-resistant Acn^D527A^, or phosphomimetic or phospho-inert versions of the conserved serine residues, S641,731, S437 (Nandi et al., 2014; Nandi et al., 2017). These studies documented elevated protein quality control autophagy for the phosphomimetic or caspase-cleavage-resistant Acn mutants compared to wild-type Acn, whereas the phospho-inert mutants failed to increase basal autophagy (Fig. 1C).

**Figure 1:**
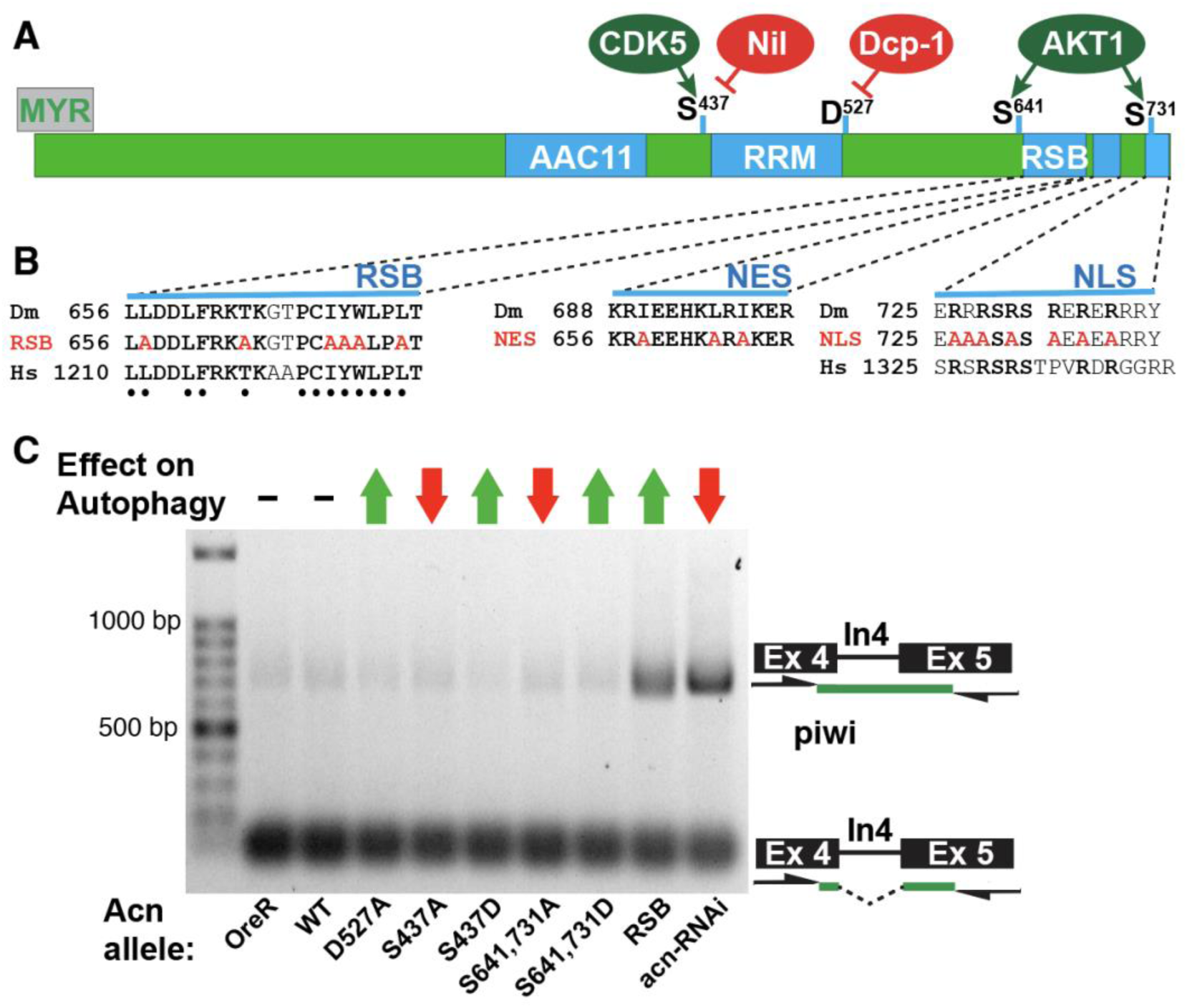
Basal autophagy regulation is independent of Acn-dependent alternative splicing. (A) Diagram depicting conserved elements of Acinus proteins, including binding site for AAC11 (Rigou et al., 2009), the RRM domain and the RSB domain interacting with and RNPS1 and Sap18 (Murachelli et al., 2012), the site of caspase-mediated cleavage (Nandi et al., 2014); and serine residues modified by AKT or CDK5-mediated phosphorylation (Hu et al., 2005; Nandi et al., 2014; Nandi et al., 2017) and Nil-mediated dephosphorylation (Nandi et al., 2022). (B) Sequences show conservation between Drosophila (Dm) and human (Hs) RSB and NLS domains, and mutations (in red) to test functional relevance of the indicated domains. Dots below the RSB sequences indicate amino acids previously identified to be critical for Acn binding to the ASAP complex (Murachelli et al., 2012). (C) RT-PCR assay reveals excision of piwi intron4 by wild-type Acinus and the indicated Acn mutants altering basal autophagy, in contrast to RNAi-mediated Acn knock down or the Acn^RSB^ mutant. Arrows on top indicate effects of different mutations that elevate (green) or reduce (red) basal autophagy as shown previously (Nandi et al., 2014; Nandi et al., 2017) or in experiments here for Acn^RSB^. Note that effects on splicing and basal autophagy are not corelated. Genotypes are listed in Suppl. Table 1.

Binding of Acn to RNPS1 and Sap18 at the RSB domain (Fig. 1A, B) has been detailed by structural studies (Murachelli et al., 2012). We mutated this previously identified RSB binding site and created the N-terminally Myc-tagged Acn^RSB^ transgene (Fig. 1 B) in the *acn* null background. The resulting flies were semi-lethal. From rare escapers, ovaries were collected to test the effect of the Acn^RSB^ mutant on splicing of intron-4 of the *piwi* gene. Consistent with previous observations that the ASAP complex is necessary for removal of intron-4 from the piwi pre-RNA (Hayashi et al., 2014; Malone et al., 2014), the Acn^RSB^ mutant failed to efficiently splice out piwi intron-4, like Acn knock down flies (Fig. 1C). By contrast, splicing of piwi intron-4 remained unaltered in other transgenic Acn mutants with increased or decreased levels of autophagy (Fig. 1C). This suggests the role of Acn in autophagy is independent of its function in alternative splicing.

To further explore a potential splicing-independent function of Acn in autophagy, we investigated localization of the splicing-compromised Acn^RSB^ protein and its effect on basal autophagy. Because gen-Acn^RSB^ flies were semilethal, we used a uas-Myc-Acn^RSB^ transgene to examine its subcellular localization compared to uas-Myc-Acn^WT^ (Fig. 2A-F). Furthermore, we tested the uas-Myc-Acn^SDR^ mutant, which in addition to the RSB mutation carries mutations (D527A and S437D) that inhibit caspase cleavage and stabilize Acn (Nandi et al., 2014; Nandi et al., 2017). Unlike Myc-Acn^WT^ (Fig. 2A), both mutant Acn variants primarily localized outside the nucleus when expressed in larval eye discs (Fig. 2B, C). Moreover, clonal expression in fat body cells marked by nuclear nGFP revealed Acn^RSB^ and Acn^SDR^ proteins enriched outside of nuclei (Fig. 2E,F), in contrast to Acn^WT^ (Fig. 2D).

**Figure 2:**
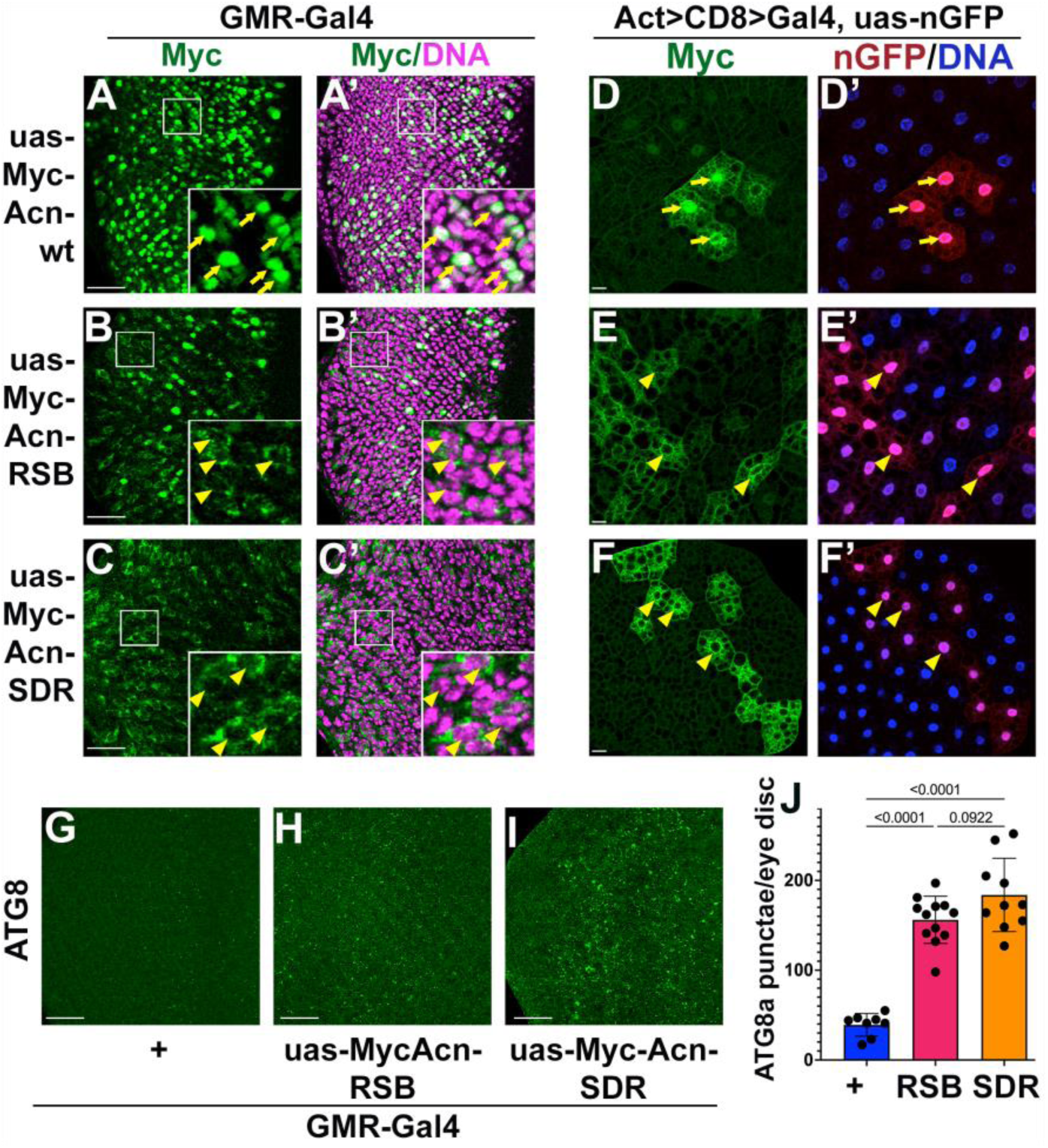
The Acn^RSB^ and Acn^SDR^ mutants are cytoplasmic and induce autophagy. (A-C) Micrographs of eye discs expressing the indicated uas-Myc-Acn transgenes under control of GMR-Gal4 stained for Myc and DNA as indicated. (D-F) Micrographs of fat bodies clonally co-expressing the indicated uas-Myc-Acn transgenes and nuclear nGFP stained for Myc and DNA as indicated. Arrows (D,D’) point to nGFP-marked nuclei positive for Myc-Acn-wt. Arrowheads point to example nuclei marked by nGFP but excluded from staining for Myc-Acn^RSB^ (E,E’) or Myc-Acn^SDR^ (F,F’). (G-I) Micrographs of eye discs expressing the indicated uas-Myc-Acn transgenes under control of GMR-Gal4 stained for ATG8. J) Quantification of ATG8 punctae in eye discs as shown in panels G-I. Bar graphs indicate means and standard deviations; significance assessed by one-way ANOVA. Scale bars are 25µm. Genotypes are listed in Suppl. Table 1.

To analyze effects on autophagy, we stained larval eye discs for ATG8a which marks both autophagosomes and early autolysosomal structures. We observed a significantly increased number of ATG8a-positive punctae in eye discs expressing uas-Myc-Acn^RSB^ or uas-Myc-Acn^SDR^ in comparison to wild-type controls (Fig. 2G-I; J).

Taken together, the Acn^RSB^ mutation interferes with Acn nuclear localization and its role in splicing but enhances Atg8-positive autophagosomes. This suggests that Acn-mediated splicing is not required for Acn-induced effects on basal autophagy and that the cytoplasmic subset of Acn promotes autophagy.

### Nucleocytoplasmic Acn transport regulates autophagy

To independently test the role of nuclear versus cytoplasmic localization for Acn in promoting autophagy and to avoid overexpression, we generated a set of genomic Acn mutants altering the nuclear to cytoplasmic ratio. Bioinformatic analysis predicted a nuclear localization signal (NLS) and a nuclear export signal (NES) in Acn (Fig. 1A). We mutated critical amino acid residues (Fig. 1B) and generated N-terminally Myc-tagged Acn mutants predicted to be enriched in cytoplasm or nucleus and expressed them in transgenes under control of the Acn promoter in an *acn* null background.

Localization of these Acn mutants was examined in eye discs and fat bodies. In 96-hr larval fat bodies, Myc-Acn^WT^ was predominantly nuclear with only a minor cytoplasmic fraction (Fig. 3A) as previously shown for endogenous Acn protein (Haberman et al., 2010). By contrast, Myc-Acn^NLS^ was largely cytoplasmic (Fig. 3B), whereas Myc-Acn^NES^ was enriched inside nuclei (Fig. 3C) lacking the small fraction of cytoplasmic Acn observed for Myc-Acn^WT^. In eye discs, Myc-Acn^WT^ was differentially expressed in different photoreceptor nuclei (Fig. 3D), consistent with dynamic cell type-specific expression of endogenous Acn protein (Haberman et al., 2010). By contrast, Myc-Acn^NLS^ showed overall reduced nuclear Acn levels (Fig. 3E). Myc-Acn^NES^ was highly enriched in all photoreceptor nuclei (Fig. 3F) and lacked the dynamic changes of wild-type Acn protein. Reduced levels of cytoplasmic Myc-Acn^NLS^ and increased levels of nuclear Myc-Acn^NES^, are consistent with susceptibility of cytoplasmic Acn to caspase-mediated degradation and the uniform expression pattern of endogenous *acn* gene (Nandi et al., 2014).

**Figure 3:**
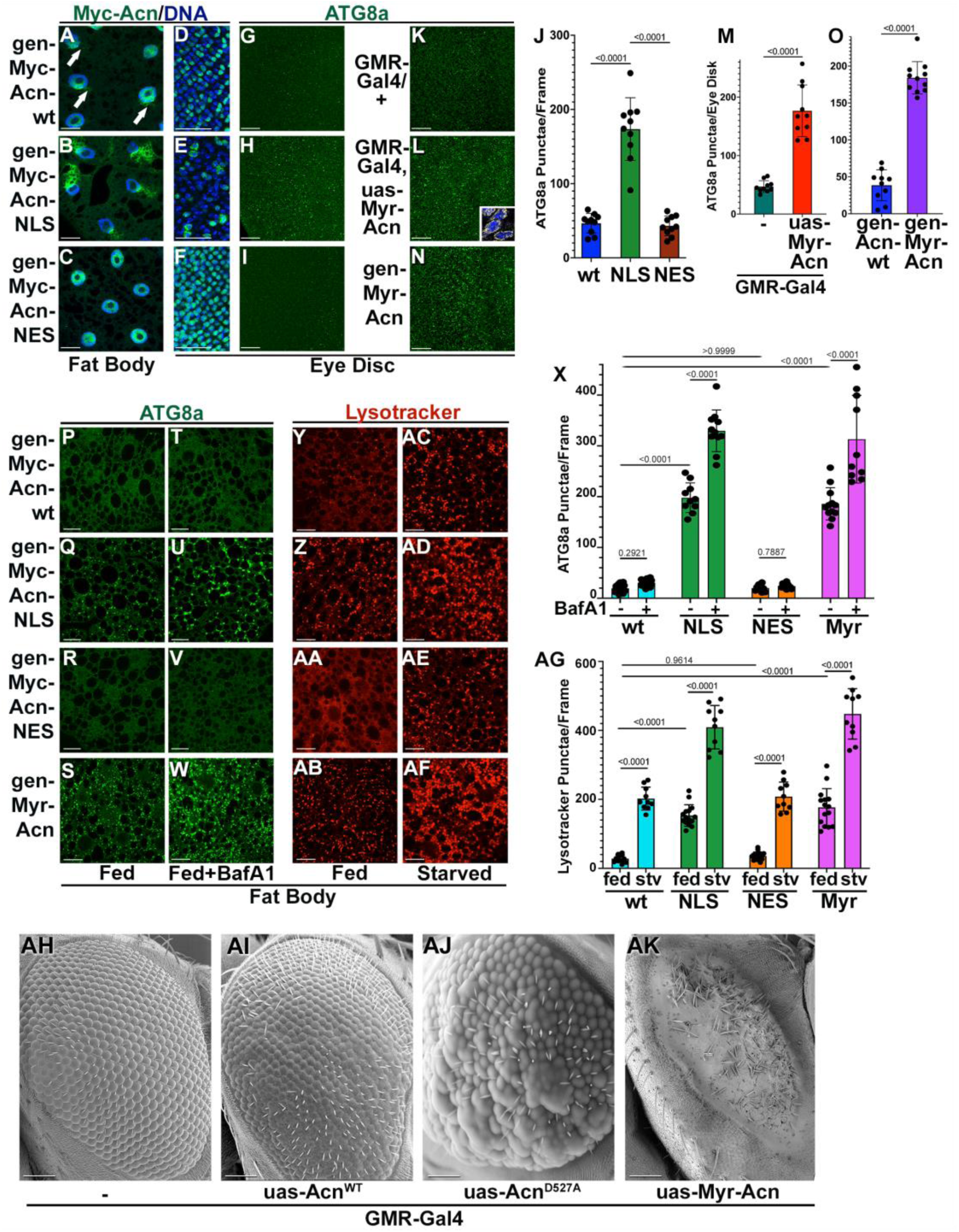
Cytoplasmic location of Acn promotes autophagy. Micrographs of fat bodies (A-C, P-AF) or eye discs at the level of photoreceptor cells (D-N) expressing the indicated genomic or uas-controlled Acn transgenes stained for Myc and DNA (A-F) or Atg8a (G-W) without (G-S) or with BafA1 treatment (T-W). (J) Quantification of ATG8 punctae in eye discs as shown in (G-I). (M) Quantification of ATG8 punctae in eye discs as shown in (K,L). (O) Quantification of ATG8 punctae in eye discs as shown in (G,N). (X) Quantification of ATG8 punctae in fat bodies treated with or without BafA1 as shown in (P-W). (Y-AF) Micrographs of lysotracker-stained fed or starved 96-h fat bodies expressing the indicated genomic Acn transgenes. (AG) Quantification of lysotracker punctae in fat bodies as shown in (Y-AF). Bar graphs show means and standard deviations; significance evaluated by one-way (J,M,O) or two-way ANOVA (X,AG). (AH-AK) SEMs of eyes from flies grown at 29°C and expressing GMR-Gal4 (AH), or in addition uas-Myc-Acn^WT^ (AI), uas-Myc-Acn^D527A^ (AJ), or uas-Myr-Acn (AK). Scale bars in A-AF are 25 µm and in AH-AK 50 µm. Genotypes are listed in Suppl. Table 1.

Next, we tested whether the altered subcellular distribution of these mutants also altered their effect on autophagy. In larval eye discs, cytoplasmic Myc-Acn^NLS^ robustly elevated basal autophagy as indicated by the increase in endogenous ATG8a punctae compared to both Myc-Acn^WT^ or Myc-Acn^NES^ (Fig. 3G-I, J). Furthermore, in well-fed 96-h fat bodies expressing Myc-Acn^NLS^, Atg8 punctae were significantly increased compared to Myc-Acn^WT^ or Myc-Acn^NES^ (Fig. 3P-R, X). To distinguish whether this increase reflects stalled autophagosomes or increased autophagy induction and flux, we used BafilomycinA1 (BafA1) to block autophagosome-lysosome fusion (Mauvezin et al., 2015). The enhancement of ATG8a punctae upon BafA1 treatment demonstrated elevated autophagic flux for Myc-Acn^NLS^, compared to wild-type Myc-Acn or nuclear-enriched Myc-Acn^NES^ (Fig. 3T-V, X). Additional support for elevated autophagic flux came from lysotracker staining to evaluate acidification of autolysosomes and lysosomes in larval fatbodies which are highly susceptible to starvation-induced autophagy (Rusten et al., 2004; Scott et al., 2004). Consistent with elevated levels of autophagic flux in Myc-Acn^NLS^ mutants, lysotracker staining was elevated in well-fed Acn^NLS^ larval fat bodies compared to Myc-Acn^WT^ or Myc-Acn^NES^ (Fig. 3Y-AA; AG). Starvation-induced mTor-regulated autophagy was unaffected by the Acn^NES^ mutation, but additive with the autophagy-enhancing effect of Myc-Acn^NLS^, consistent with Acn and mTOR regulating autophagy by independent pathways (Fig. 3AC-AE, AG).

Taken together, these data indicate that NLS and NES sequences direct the subcellular distribution of Acn proteins between nucleus and cytoplasm with the cytoplasmic fraction of Acn enhancing basal autophagy by a pathway that is independent of mTOR inhibition.

### Membrane-targeted myristoylated Acn promotes autophagy

To further test the role of cytoplasmic localization for Acn, we added an N-terminal myristoylation signal to generate Myr-Acn transgenes (Fig. 1A). Expression of uas-Myr-Acn under control of GMR-Gal4 yielded membrane-enriched Acn (inset Fig. 3L) which caused significantly elevated ATG8 staining in eye discs compared to wild-type controls (Fig. 3K,L, M).

For further tests, we generated a genomic Myr-Acn transgene under control of the endogenous Acn promoter. Eye discs expressing gen-Myr-Acn exhibited a significant increase in ATG8a-positive punctae compared to wild-type controls, consistent with elevated levels of autophagy (Figure 3G,N, O). Similarly, 96h-old fat bodies of fed gen-Myr-Acn larvae demonstrated elevated numbers of Atg8a punctae in comparison to wild-type controls (Fig. 3P,S; X). To assess autophagic flux, we stained fat bodies for Atg8a after BafA1 treatment. We observed significantly increased autophagic flux upon expression of genomic Myr-Acn (Fig. 3S,W; X).

Furthermore, under fed conditions, the number of lysotracker-positive punctae in gen-Myr-Acn was significantly elevated compared to wild-type controls (Fig. 3Y,AB; AG). For both genotypes, starvation further increased lysotracker staining (Fig. 3Y,AB,AC,AF; AG), consistent with additive effects for the different pathways regulated by either Acn or mTOR.

GMR-Gal4-driven elevated Acn disrupts eye development due to increased Atg1 kinase activity (Tyra et al., 2020). The resulting "rough-eye" phenotype (Fig. 3AH,AI) was more severe when Acn activity was increased by Cdk5-mediated phosphorylation or a corresponding phosphomimetic mutation (Nandi et al., 2017), or by blockage of its caspase-mediated cleavage in the Acn^D527A^ mutation (Fig. 3AJ). Myristoylated Acn further worsens the eye phenotype with dramatically reduced eye size (Fig. 3AK), suggesting that membrane targeting further enhances Acn function and thereby Atg1 activity.

### BioID approach links Acn to Rab32

To identify potential interactors that may act downstream of cytoplasmic Acn, we used a BioID approach. We fused the microID2 (µID) derivative of BirA (Johnson et al., 2022) to the N-terminus of cytoplasmically-enriched Acn^SDR^ and nuclear Acn^NES^ and generated corresponding Quas-controlled transgenes (Fig. 4A). To avoid developmental consequences of elevated Acn activity (Tyra et al., 2020), we restricted expression to adult photoreceptors using the CarT-QF2 driver (Shekhar et al., 2023). Western blots of head lysates revealed several biotinylated proteins specific to flies expressing the µID-tagged Acn proteins (Fig. 4B).

**Figure 4:**
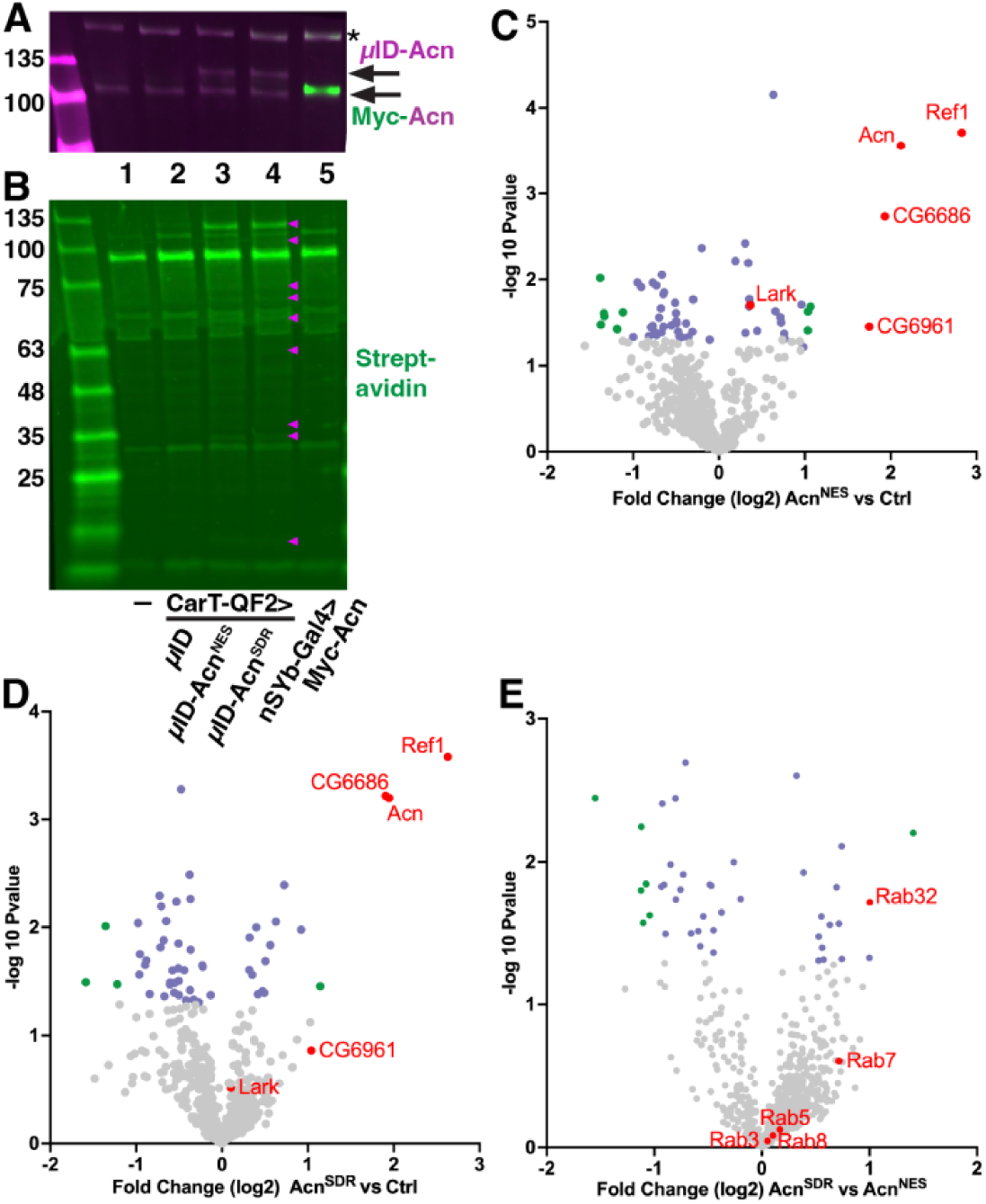
µID-Acn^SDR^ and µID-Acn^NES^ detect Acn associated proteomes in adult photoreceptors. (A,B) Western blots of head lysates from wild-type flies (lane 1) or flies expressing under control of the CarT-QF2 driver Quas-µID (2), Quas-µID-Acn^NES^ (lane 3), Quas-µID-Acn^SDR^ (lane 3), or flies expressing uas-Myc-Acn under control of the nSyB-Gal4 driver (lane 5) to show size of Acn lacking the µID fusion. (A) Proteins were identified with antibodies against (A) Acn (magenta) or Myc (green). The star indicates a non-specific band. (B) Biotin containing proteins were identified using streptavidin. Arrowheads point to bands specific to lanes with µID-tagged Acn proteins. (C,D) Volcano plots showing significantly enriched biotinylated proteins in µID-Acn^NES^ (C) or µID-Acn^SDR^ (D) compared to control µID-Acn. Enriched proteins related to RNA metabolism are highlighted in red. (E) Volcano plot showing significantly enriched proteins in µID-Acn^SDR^ compared to µID-Acn^NES^. Rab32 and other detected Rab proteins are highlighted in red. For all volcano plots, proteins with p < 0.05 & lFC > 2 appear in green, p < 0.05 & FC < 2 in purple, and not significant in grey, except groups of proteins highlighted in red. Source data for volcano blots are in Supplemental Table 2.

For analysis of the Acn-associated proteomes by mass spectroscopy, heads were harvested and biotinylated proteins enriched (Uçkun et al., 2021). Compared to control samples, biotinylated proteins from heads expressing µID-Acn^NES^ (Fig. 4C) or µID-Acn^SDR^ (Fig. 4D) were both enriched for multiple RNA-interacting proteins, consistent with the role of Acn in RNA metabolism. These included the RNA export factor binding protein1, Ref1 (Lasko, 2000), CG6686, the ortholog to the splicing-related protein Sart1 (Laetsch et al., 2014), the RNA-binding protein CG6961 (Lasko, 2000), and also Acn itself (Fig. 4C,D).

A different picture emerged from the direct comparison of the proteomes associated with µID-Acn^SDR^ relative to µID-Acn^NES^ (Fig. 4E). Consistent with cytoplasmic Acn regulating membrane trafficking, several Rab proteins appeared to be slightly enriched in the µID-Acn^SDR^ samples. Enrichment was strongest for Rab32, a small GTPase previously connected to autophagy (Wang et al., 2011). More recently, the Rab32^S71R^ mutant was identified as a risk factor for Parkinson’s disease (Gustavsson et al., 2024; Hop et al., 2024). This disease link for Rab32 was strengthened by its function in the membrane recruitment of the Lrrk2 kinase (Gustavsson et al., 2024; McGrath et al., 2021; Waschbüsch et al., 2014) as pathogenic variants of Lrrk2 are linked to Parkinson’s disease as well (reviewed in Bentley-DeSousa et al., 2025a).

Support for a role of Rab32 and Lrrk in Acn activity came from genetic interactions. Compared to wild-type eyes (Fig. 3AH), GMR-Gal4-driven expression of Myr-Acn during eye development yielded nearly complete loss of adult eyes (Fig. 3AK, 5A). This effect was partially suppressed by loss of one allele of *Rab32* (Fig. 5B) or *Lrrk* (Fig. 5C) yielding rough but recognizable eyes (quantified in Fig. 5U). *Rab32*^ltd-1^ is a loss-of-function allele deleting the C-terminal 203 amino acids shared by all splice variants (Ma et al., 2004); *lrrk*^ex1^ is a loss-of-function allele that deletes 464 bp of the C-terminal coding sequence including the WD40 domain and part of the kinase domain (Lee et al., 2007).

**Figure 5:**
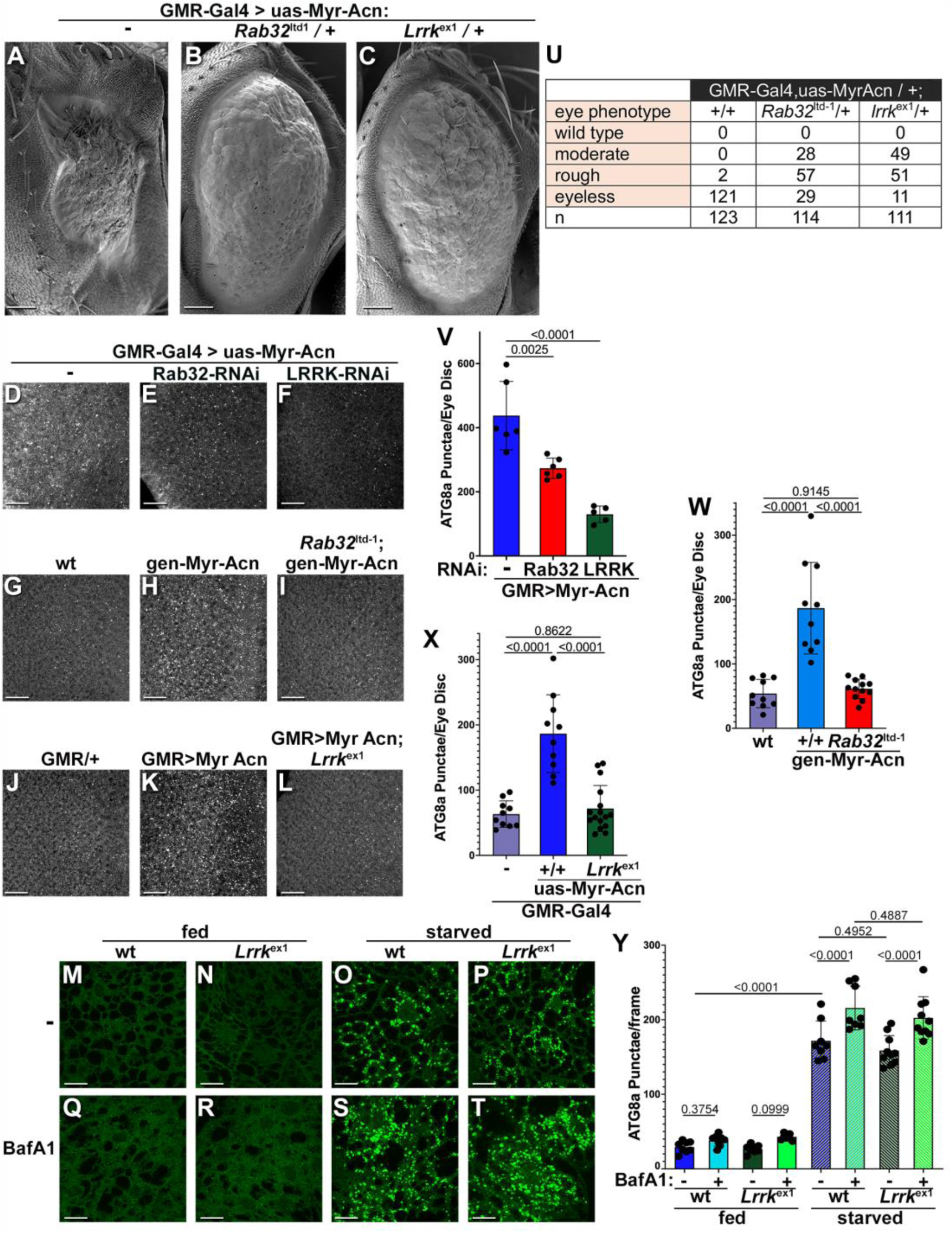
Lrrk is required for Acn-induced, but not starvation-induced autophagy. (A-C) SEMs of eyes from flies expressing uas-Myr-Acn under GMR-Gal4 control at 29°C (A), and in addition are heterozygous for *Rab32*^ltd-1^(B) or *Lrrk*^ex1^ (C). Quantification of eye phenotypes in (U). (D-F) Micrographs of eye discs expressing uas-Myr-Acn under GMR-Gal4 control (D) and in addition uas-Rab32-RNAi (E) or uas-Lrrk-RNAi (F) stained for Atg8a. Quantification of Atg8a punctae in (V). (G-I) Micrographs of wt eye discs (G) or eye discs expressing gen-Myr-Acn in otherwise wild-type (H) or *Rab32*^ltd-1^ (I) background stained for Atg8a. Quantification of Atg8a punctae in (W). (J-L) Micrographs of eye discs expressing GMR-Gal4 (J) and uas-Myr-Acn in otherwise wild-type (K) or *Lrrk*^ex1^ (L) background stained for Atg8a. Quantification of Atg8a punctae in (X). (M-T) wild-type (M,O,Q,S) or *Lrrk*^ex1^(N,P,R,T) 96-h fat bodies untreated (M-P) or BafA1-treated (Q-T) and either fed (M,N,Q,R) or starved (O,P,S,T) stained for Atg8a. Quantification of Atg8a punctae in fat bodies as shown in (Y). (U-Y) Bar graphs show means and standard deviations with significance assessed by one-way (V-X) or two-way ANOVA (Y). Scale bars in A-C are 50 µm; D-T are 25 µm. Genotypes are listed in Suppl. Table 1.

We further tested a role of *Rab32* and *Lrrk* downstream of Acn in autophagy. To induce Acn-activated autophagy, we expressed uas-Myr-Acn under control of GMR-Gal4 in eye discs (Fig. 5D). Levels of Atg8a punctae were significantly reduced by co-expressed RNAi transgenes to knock down Rab32 (Fig. 5E,V) or Lrrk (Fig. 5F,V). Furthermore, autophagy activated by gen-Myr-Acn (Fig. 3S,AG) was completely suppressed in *Rab32*^ltd-1^ mutant eye discs (Fig. 5G-I,W). Similarly, the *Lrrk*^ex1^ loss-of-function allele suppressed autophagy induction by GMR-Gal4-driven uas-Myr-Acn in eye discs (Fig. 5J-L,X). Together these data argue for a requirement of Rab32 and Lrrk in Acn-activated autophagy.

Importantly, this does not reflect a global requirement of *Lrrk* for autophagy. In contrast to Acn-activated basal autophagy, starvation-induced mTOR-regulated autophagy was unaffected by loss of *Lrrk* function (Fig. 5M-T; quantified in 5Y). Our findings are consistent with a specific requirement for *Lrrk* in Acn-regulated protein quality control autophagy, but not in mTOR-regulated starvation-induced autophagy. The findings add further support for Acn activating autophagy by a distinct pathway that is independent of mTOR inhibition.

## Discussion

Autophagy is critical for cellular homeostasis with numerous signaling pathways impinging on its regulation. Here, we have focus on one of these pathways that is regulated by the activity of Acn. Acn-dependent autophagy is engaged by expression of aggregation-prone proteins like polyQ proteins or Aβ42 peptides, and activated by CDK5-mediated Acn phosphorylation (Nandi et al., 2017). Downstream of activated Acn, an unknown mechanism activates Atg1 (Tyra et al., 2020), the master regulatory kinase of autophagy. Acn and mammalian Acin1 orthologues are mostly nuclear proteins with well-established roles in alternative splicing (Fukumura et al., 2024; Hayashi et al., 2014; Malone et al., 2014; Murachelli et al., 2012; Schwerk et al., 2003), which originally suggested an indirect, nuclear role for Acn in regulating autophagy.

Surprisingly, we find that the splicing-related function of nuclear Acn, based on its involvement in the ASAP complex, is not required for autophagy induction by Acn. This conclusion is based on several different Acn mutants that either disrupt its binding to the ASAP complex (Acn^RSB^), interfere with its normal nucleocytoplasmic shuttling (Acn^NLS^, Acn^NES^), or target Acn to membranes (Myr-Acn). Together, our analysis of these Acn mutants points to a critical function of the cytoplasmic fraction of Acn in promoting autophagy.

Consistent with a direct role of cytoplasmic Acn in membrane trafficking, proximity labeling of the associated proteome identified enrichment for several Rab proteins. Among them, only Rab32 was significantly enriched, and genetic analysis substantiated its requirement in Acn-induced autophagy. Rab32 has been linked to multiple roles in membrane trafficking. In *Drosophila,* the classic *Rab32*^ltd-1^ allele was identified as mutant disrupting pigment granule formation (Lloyd et al., 1998; Ma et al., 2004), in line with defects in lysosome-related organelles associated with mammalian Rab32 mutants (Bowman et al., 2019). Furthermore, consistent with our findings, Rab32 contributes to the formation of autophagosomes under basal conditions in *Drosophila* and mammalian cells (Hirota and Tanaka, 2009; Wang et al., 2011).

More recently, Rab32 has attracted attention from two interrelated observations: the identification of the Rab32^S71R^ mutant as a risk factor for Parkinson’s disease (Gustavsson et al., 2024; Hop et al., 2024) and its role in membrane recruitment of Lrrk2 (Gustavsson et al., 2024; McGrath et al., 2021; Waschbüsch et al., 2014). Importantly, multiple Lrrk2 mutations are linked to Parkinson’s disease as well (reviewed in Bentley-DeSousa et al., 2025a). This set of observations and previous work showing rescue of *Drosophila lrrk* phenotypes by expression of human Lrrk2 (Dodson et al., 2014; Soukup et al., 2016) prompted our experiments revealing the critical role of Lrrk in autophagy induction by activated Acn.

A large and complicated literature connects Lrrk2 to aspects of autophagy and lysosomal delivery, with several studies suggesting an inhibitory role of Lrrk2 in autophagy (Berth et al., 2022; Madureira et al., 2020). Much of recent work has focused on Lrrk2 recruitment during conjugation of ATG8 to single membranes (CASM), a common response to disruption of the pH balance of different acidified organelles (Durgan and Florey, 2022) with possible linkage to Parkinson’s disease (Bentley-DeSousa et al., 2025b; Eguchi et al., 2024; Kuwahara and Iwatsubo, 2024; Sakurai and Kuwahara, 2025). Characteristics of CASM are, however, not consistent with our observations of Acn-activated autophagy. Most importantly, BafA1 inhibits membrane ATG8ylation during CASM (Durgan and Florey, 2022). By contrast, robust Atg8a staining with activated Acn was further enhanced by BafA1 treatment, consistent with elevated autophagic flux. Furthermore, Acn acts through ATG1/ULK1 activation (Tyra et al., 2020), whereas CASM is independent of ATG1/ ULK1 activation (Durgan and Florey, 2022). Together these data indicate that the requirement of Lrrk for Acn-induced autophagy is independent of its role in CASM.

A separate line of evidence connects the Lrrk2 kinase to autophagy and lysosomal delivery through its membrane recruitment by Rab32, or the related Rab29 and Rab38 GTPases (McGrath et al., 2021) and its role in phosphorylating multiple Rab proteins (Pfeffer, 2026; Steger et al., 2016). These phosphorylation events can switch effector proteins preferentially binding to GTP-bound Rab proteins with consequences for trafficking of endolysosomal and secretory compartments (Pfeffer, 2023). The diversity of Rab proteins expressed and phosphorylated in different cell types and the speed of their dephosphorylation (Berndsen et al., 2019; Dou et al., 2023; Fricke et al., 2026; Pfeffer, 2026) may account for the variability of consequences in response to loss of Lrrk2 function in different physiological contexts (Dodson et al., 2014; Dou et al., 2023; Soukup et al., 2016; Tong et al., 2012). Seemingly contradictory effects of Lrrk2 inhibiting or activating autophagy may also reflect the diversity of substrates other than Rab proteins, including EndophilinA and Beclin (Berth et al., 2022; Madureira et al., 2020; Manzoni et al., 2016; Soukup et al., 2016).

Our data strongly support the importance of the Lrrk and Rab32 downstream of activated Acn for the induction of autophagy, but our study has several limitations. We have not been able to detect direct interactions between Acn and either Rab32 or Lrrk suggesting that additional intermediates could mediate their interaction. Similarly, it remains to be resolved whether the Lrrk and Atg1/ULK1 kinases promote autophagy downstream of activated Acn independently of each other or through the same pathway. Intriguingly, mTOR-independent elevation of ULK1 phosphorylation has been observed in response to prolonged inhibition of Lrrk2 kinase activity (Manzoni et al., 2018) suggesting an indirect interaction between the two kinases. Furthermore, both kinases have been implicated in axonal transport of autophagosomes, but by distinct mechanisms with effects on kinesin or dynein-mediated transport (Dou et al., 2023 ; Fricke et al., 2026; Madureira et al., 2020; Toda et al., 2008). Whether activated Acn similarly affects axonal transport remains to be tested.

## Materials and Methods

### Key resources table

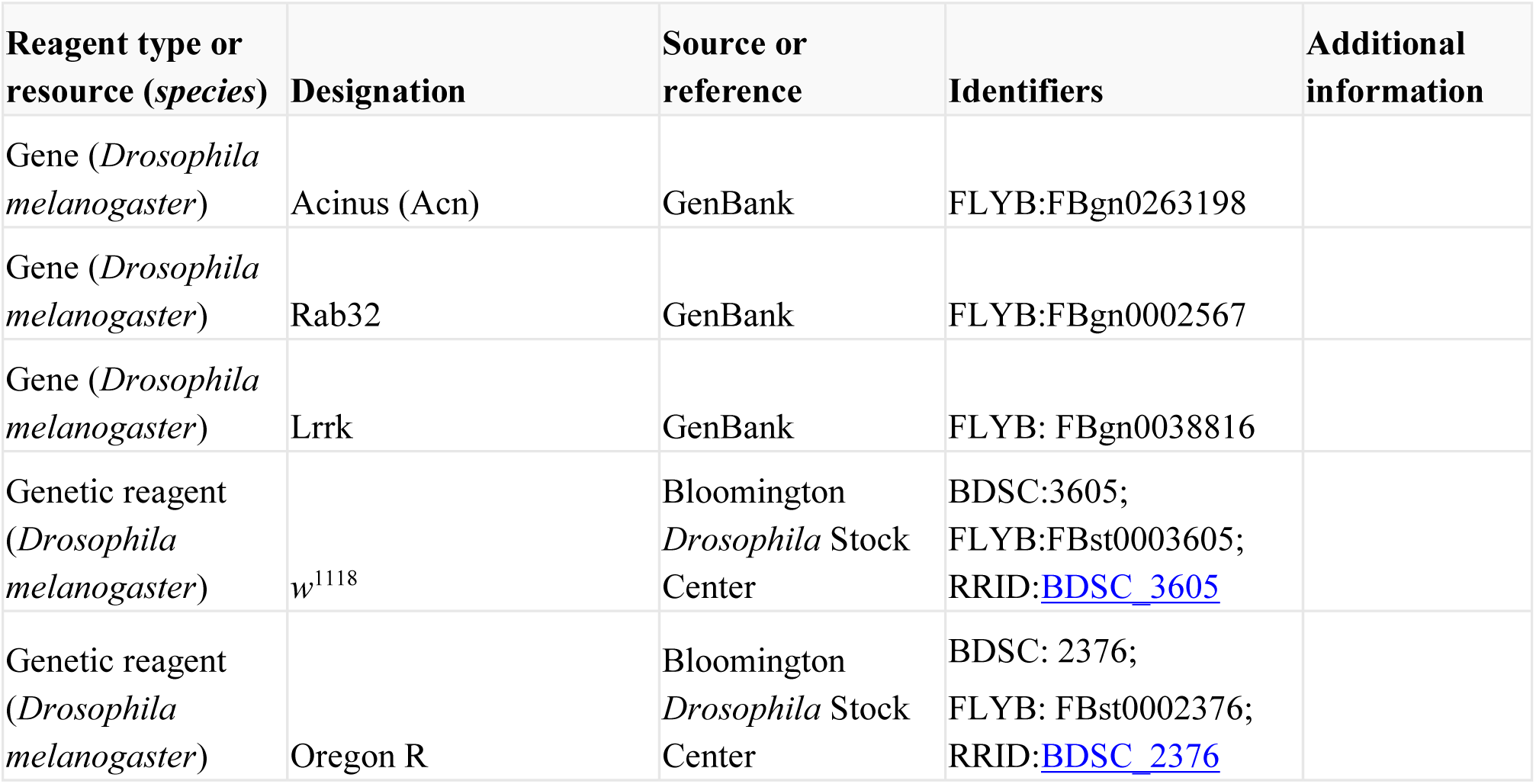
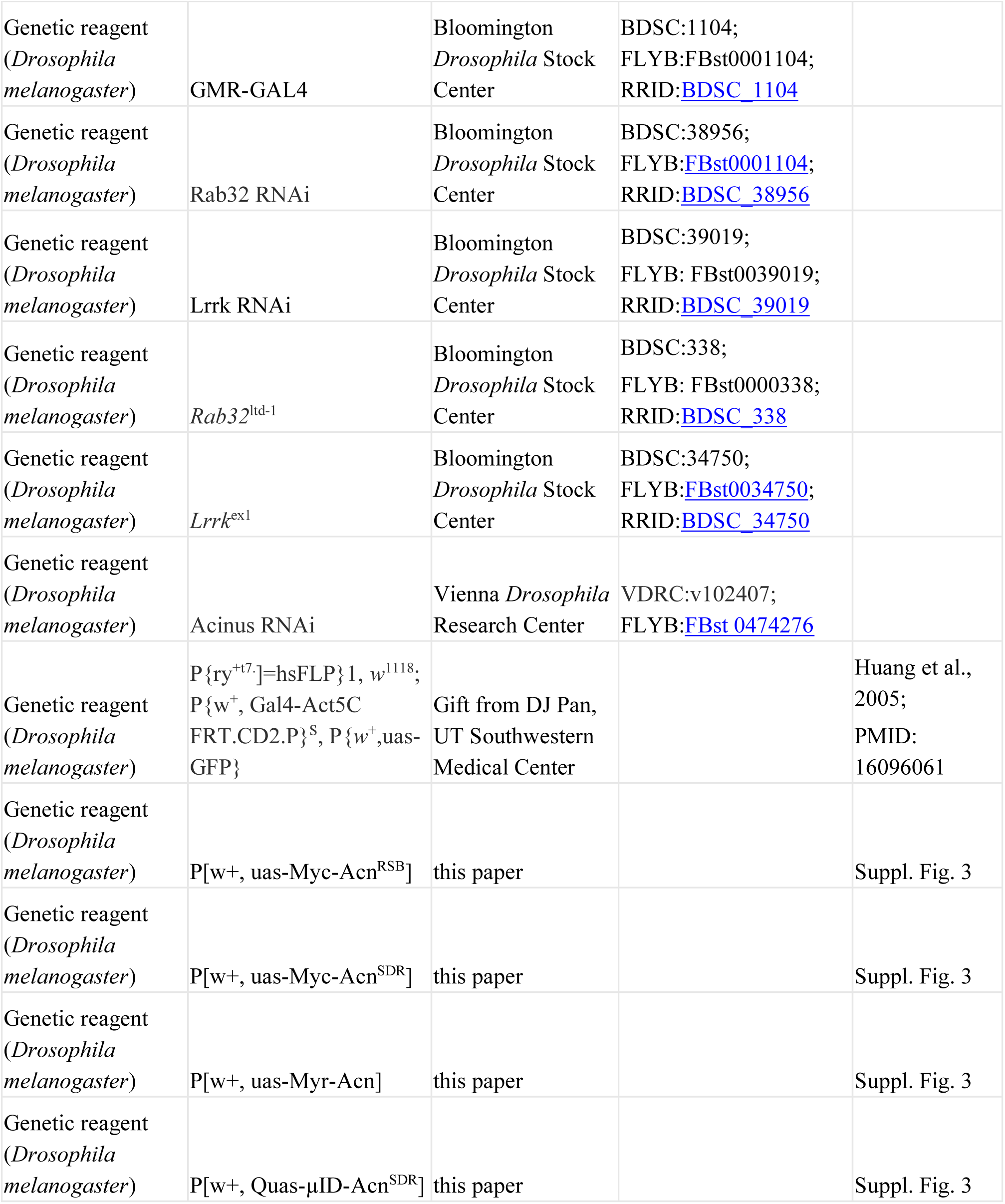
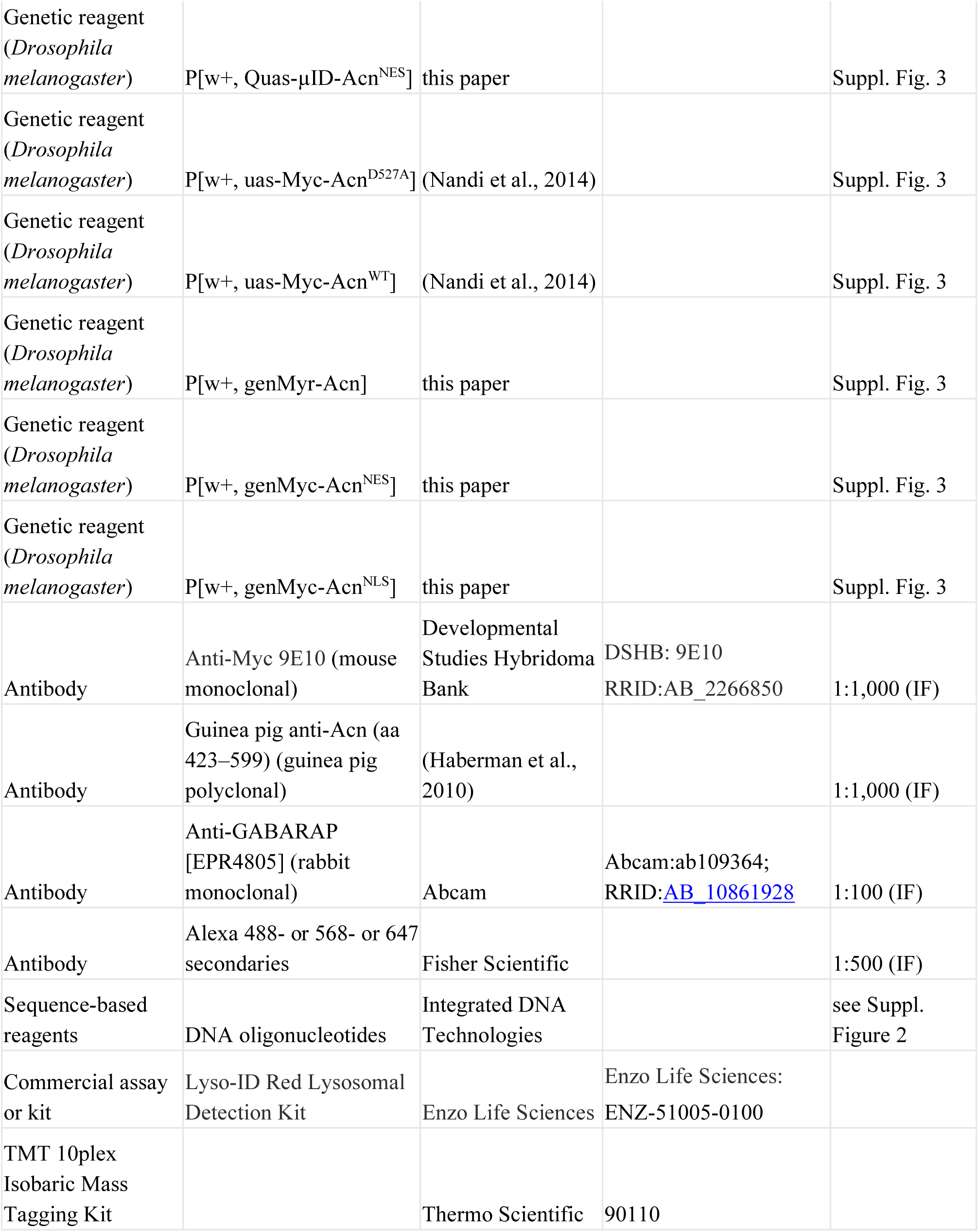
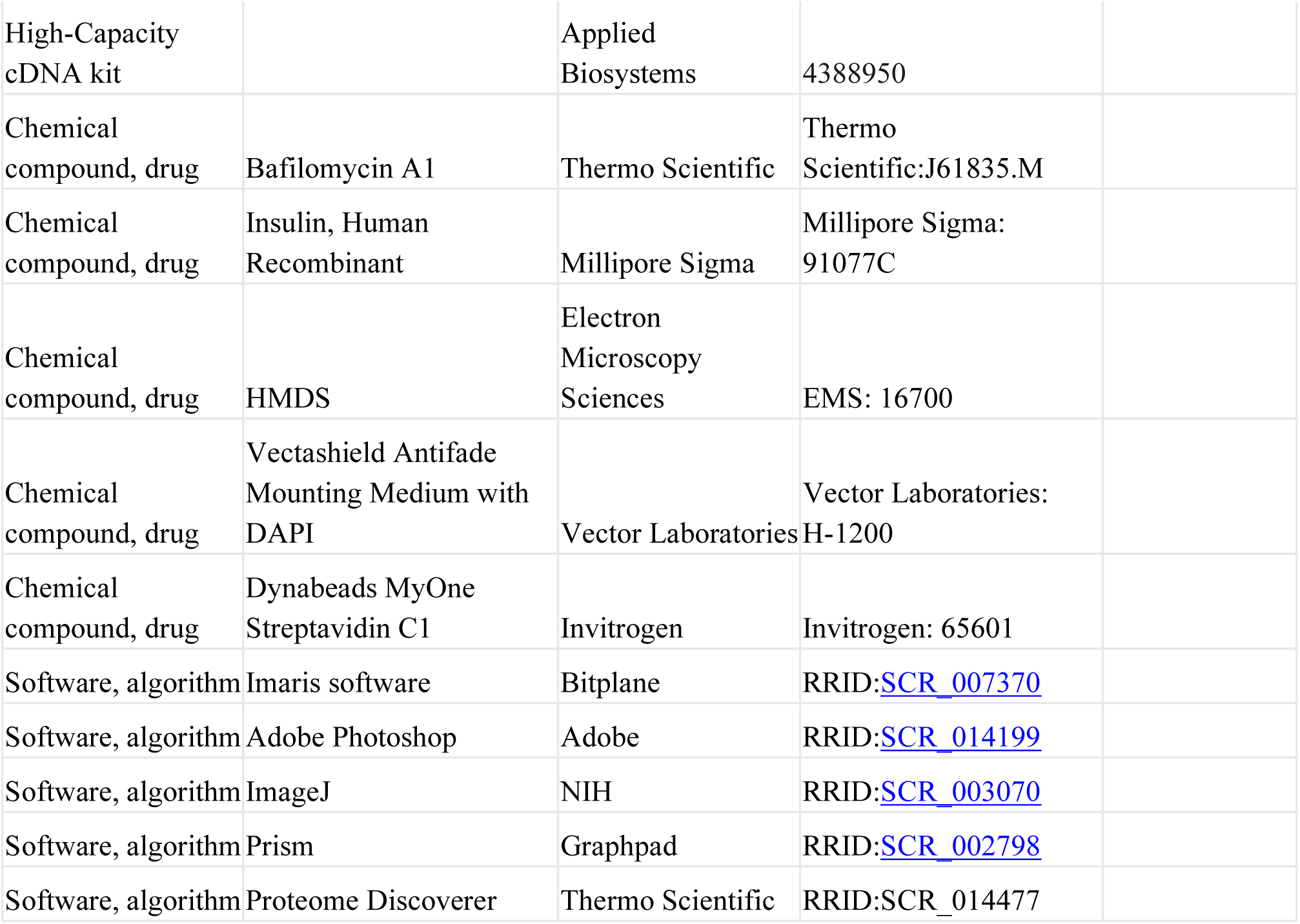

### Experimental model

Fly stocks were maintained at room temperature under standard conditions. To maximize knockdown efficiency, experiments with uas-RNAi transgenes were performed at 29°C. Overexpression clones of uas-Acn transgenes in fat bodies were generated as described (Tyra et al., 2020) after crossing to hsFLP; Actin>CD2>Gal4, uas-GFP and exposing 3-day old larvae to a heat shock for 15 min at 38°C

Drosophila lines used in this study were provided by the Bloomington *Drosophila* Stock Center, Vienna Drosophila Research Center, gifted by DJ Pan or generated in this lab as listed in the Resources Table.

BestGene, Inc injected DNA for generation of transgenic flies. A 4-kb *Acn* genomic fragment previously shown to be sufficient for genomic rescue (Haberman et al., 2010) was modified by site-directed mutagenesis to generate Acn mutants under control of its endogenous promoter. The resulting constructs were sequence-verified, cloned into an AttB vector, and integrated into the AttP landing site at 96F3 (Venken et al., 2006). Similarly, uas-driven wild-type and mutant Acn transgenes were generated by standard mutagenesis from full-length Acn cDNA, verified by sequencing, and cloned into pAttB-uas vector as described (Nandi et al., 2017). For the generation of Quas-µID-Acn transgenes, the µID encoding DNA (Johnson et al., 2022) was generated as gBlock (IDT) and cloned into the pAttb-Quas vector, N-terminal to Acn^NES^ or Acn^SDR^ or a stop codon for controls. For complete sequences see Supplemental Figure 2. All uas and Quas controlled Acn transgenes were inserted into the 43A1 landing site (Venken et al., 2006).

### Histology

Scanning Electron Micrographs (SEMs) of fly eyes were obtained as described (Wolff, 2011). Briefly, eyes were fixed for 2 hr in 2% paraformaldehyde, 2% glutaraldehyde, 0.2% Tween 20, and 0.1 M cacodylate buffer (pH 7.4). Subsequently, after four 12-h washes in a graded ethanol series from 25–100%, the samples were subjected to a graded hexamethyldisilazane (HMDS) series 25–100% in ethanol, each step lasting one hour. After overnight drying, flies were mounted on SEM stubs and, restricted to their bodies coated in fast-drying silver paint. Flies were sputter-coated with a gold/pallidum mixture for 90 s and imaged at 130 × magnification, with extra-high tension set at 3.0 kV on an InLens detector-equipped scanning electron microscope (SIGMA; Carl Zeiss).

### Immunofluorescence

Whole-mount tissues were processed for immunofluorescence staining as previously described (Nandi et al., 2014). For amino-acid starvation, larvae were fed a 20% sucrose solution for 3 hr. Larval eye discs or fat bodies dissected from fed or starved size-matched 90–96 hr larvae were fixed in periodate-lysine-paraformaldehyde, washed in PBS, permeabilized with 0.3% saponin in PBS (PBSS), and blocked with 5% goat serum in PBSS. Samples were then incubated with the indicated primary and secondary antibodies. Samples mounted in Vectashield containing DAPI (Vector Laboratories) were imaged using an inverted confocal microscope (LSM 710; Carl Zeiss, Jena, Germany) equipped with 63×/1.4 NA or 40×/1.3 NA Plan-Apochromat objectives. Confocal Z-stacks of eye discs and fat bodies were acquired at 1-µm intervals.

For analyzing autophagy flux, dissected fat body tissues were incubated ex vivo for 6 h with 200 nM BafA1. For assessment of basal autophagy, 10 μg ml^−1^ insulin (Sigma) was added during the BafA1 treatment but omitted for measurement of starvation-induced autophagy (Mauvezin et al., 2015).

For lysotracker staining, fat bodies were dissected from fed and starved size-matched larvae at 90–96 hr and stained with the GFP-Certified Lyso-ID Red Lysosomal Detection Kit (Enzo Life Sciences), as previously described (Rusten et al., 2004; Scott et al., 2004). Briefly, larvae were dissected in Schneider’s Drosophila medium (Gibco), the carcasses inverted to expose the fat bodies and incubated in 100 µM lysotracker Red DND-99 for 1 min. The inverted carcasses were washed with 1× PBS, and mounted on a drop of Vectashield (Vector Laboratories). Samples were imaged immediately using an inverted confocal microscope (LSM 710; Zeiss) equipped with a 63×/1.4 NA Plan-Apochromat objective. Single optical sections of the larval fat bodies were acquired for analysis and lysotracker-positive punctae were quantified using Imaris software (Bitplane).

For quantification of Atg8a puncta, Z-projections of three optical sections of larval fat body and eight optical sections of larval eye disc acquired at 1 µm intervals were analyzed using Imaris software (Bitplane). The number of lysotracker and Atg8a puncta was quantified per 1024 × 1024-pixel frame of larval fat body tissue. Images prepared for presentation were imported into Photoshop (Adobe, RRID:SCR_014199) and adjusted for gain, contrast, and gamma.

### Head collection

For Western blots, 10 dissected fly heads were collected and pestle-ground in SDS loading buffer. 0.5 head were loaded per lane. Primary antibodies (guinea pig anti-Acn and anti-Myc 9E10) were detected with IR-dye labeled secondary antibodies using the Odyssey scanner (LI-COR Biosciences). Pre-stained molecular weight markers were obtained from UBP-Bio.

For mass spectrometry experiments, flies were frozen at -80°C and heads prepared by shaking and sieving as described (Heim et al., 2022). Triplicates of approximately 0.5 ml of heads were collected for each sample and stored at -80°C until use. Heads were pestle-ground in RIPA buffer at 4°C, debris removed by centrifugation (20 min at 14,000 rpm) and biotinylated proteins collected on 20 µl Dynabeads™ MyOne™ Streptavidin C1 magnetic beads for mass spectrometry as described (Uçkun et al., 2022).

### Mass spectrometry

Mass spectrometry of proteins bound to Streptavidin beads was performed by the UT Southwestern Proteomics Core. In brief, proteins on bead were trypsin digested and released peptides labeled with the TMT 10plex Isobaric Mass Tagging Kit (Thermo). Combined sample were analyzed on a Thermo Orbitrap Eclipse MS system coupled to an Ultimate 3000 RSLC-Nano liquid chromatography system. Real-time search was used for selection of peaks for SPS-MS3 analysis, with searched performed against the Drosophila melanogaster reviewed protein database from UniProt (downloaded Sept. 20, 2021, 21173 entries). Up to 1 missed tryptic cleavage was allowed, with carbamidomethylation (+57.0215) of cysteine and TMT reagent (+229.1629) of lysine and peptide N-termini used as static modifications and oxidation (+15.9949) of methionine used as a variable modification. MS3 data were collected for up to 10 MS2 peaks which matched to fragments from the real-time peptide search identification, in the orbitrap at a resolution of 50,000, HCD collision energy of 65% and a scan range of 100–500. Protein identification and quantification were done using Proteome Discoverer v.3.0 SP1 (Thermo). Reporter ion intensities were reported, with further normalization performed by using the total intensity in each channel to correct discrepancies in sample amount in each channel. The false-discovery rate (FDR) cutoff was 1% for all peptides.

### Piwi RT-PCR

Flies were OreR or *acn* null (*acn*^1^/*acn*^27^) with transgenes inserted at 96F expressing different Acn mutants under control of the acn promoter or flises . Ovaries (5-10) were dissected from adult flies that were fed on yeast paste for at least 2 days and homogenized in Trizol (Ambion) and RNA was extracted as suggested by the manufacturer. cDNA was synthesized from 2μg RNA (in 20μL) using a High-Capacity cDNA kit (Applied Biosystems) according as suggested by the manufacturer. After PCR amplification (for primers see Supplemental Figure 3) reaction products from spliced (81 bp) and unspliced intron 4 (685 bps) were separated by standard agarose gel electrophoresis.

### Statistical methods

Statistical significance was determined in Prism software (Graphpad, RRID:SCR_002798) using one-way analysis of variance (ANOVA) for multiple comparisons, followed by Tukey’s test. We used two-way ANOVA for multiple comparisons, followed by Tukey’s test for individual comparisons to separate effects of treatment and genetic background. Bar graphs generated from this analysis demonstrate means ± SD. p values smaller than 0.05 are considered significant, and values are indicated on the graphs.

## Acknowledgements

We thank Drs. Shashank Shekhar and Ranjan Sahu for comments on the manuscript. We thank the Bloomington Stock Center (NIH P40OD018537) for flies, the Developmental Studies Hybridoma Bank at The University of Iowa for antibodies, for SEM imaging the UT Southwestern Electron Microscopy Core Facility (supported by NIH 1S10OD020103-01) and the UT Southwestern Proteomics core for mass spectrometry and data analysis. The work was supported by the NIH [R01EY10199, R01EY033184 to H.K.]

