## Supplemental files for "Autophagy induction by cytoplasmic Acinus requires Lrrk activity"

### Supplementary File 1. Genotypes of Flies used in each Figure

| **Figure 1** | **Genotype** |
|---|---|
| 1 C | OreR<br>$w^{*}$ ; *acn*$^{27}$; P[$w^{+}$, *acn*P-Myc-Acn$^{WT}$]$^{96F3}$<br>$w^{*}$ ; *acn*$^{27}$; P[$w^{+}$, *acn*P-Myc-Acn$^{D527A}$]$^{96F3}$<br>$w^{*}$ ; *acn*$^{27}$; P[$w^{+}$, *acn*P-Myc-Acn$^{S437A}$]$^{96F3}$<br>$w^{*}$ ; *acn*$^{27}$; P[$w^{+}$, *acn*P-Myc-Acn$^{S437D}$]$^{96F3}$<br>$w^{*}$ ; *acn*$^{27}$; P[$w^{+}$, *acn*P-Myc-Acn$^{S641,731A}$]$^{96F3}$<br>$w^{*}$ ; *acn*$^{27}$; P[$w^{+}$, *acn*P-Myc-Acn$^{S641,731D}$]$^{96F3}$<br>$w^{*}$ ; *acn*$^{27}$; P[$w^{+}$, *acn*P-Myc-Acn$^{RSB}$]$^{96F3}$<br>w* ; CyO / uas-Acn-RNAi$^{(KK111555)v102407}$; Da-Gal4 / + |
| **Figure 2** | **Genotype** |
| 2 A, A’ | $w^{*}$ ; GMR-Gal4/ P[w+, uas-myc-Acn$^{WT}$] $^{43A1}$; *Sb* / + |
| 2 B, B’ | $w^{*}$ ; GMR-Gal4/ P[w+, uas-myc-Acn$^{RSB}$] $^{43A1}$; *Sb* / + |
| 2 C, C’ | $w^{*}$ ; GMR-Gal4/ P[w+, uas-myc-Acn$^{SDR}$] $^{43A1}$; *Sb* / + |
| 2 D, D’ | *w**, hsFLP/+; +/+; P[w+, Gal4-Act5C (FRT.CD2).P]$^{S}$, P[w+, uas-GFP]/P[w+,uas-myc-Acn$^{WT}$]$^{43A1}$ |
| 2 E, E’ | *w**, hsFLP/+; +/+; P[w+, Gal4-Act5C (FRT.CD2).P]$^{S}$, P[w+, uas-GFP]/P[w+,uas-myc-Acn$^{RSB}$]$^{43A1}$ |
| 2 F, F’ | *w**, hsFLP/+;+/+; P[w+, Gal4-Act5C (FRT.CD2).P]$^{S}$, P[w+, uas-GFP]/P[w+, uas-myc-Acn$^{SDR}$]$^{43A1}$ |
| 2 G | $w^{*}$ ; GMR-Gal4 / +; *Sb* / + |
| 2 H | $w^{*}$ ; GMR-Gal4/ P[w+, uas-myc-Acn$^{RSB}$] $^{43A1}$; *Sb* / + |
| 2 I | $w^{*}$ ; GMR-Gal4/ P[w+, uas-myc-Acn$^{SDR}$] $^{43A1}$; *Sb* / + |
| 2 J | $w^{*}$ ; GMR-Gal4 / +; *Sb* / +<br>$w^{*}$ ; GMR-Gal4/ P[w+, uas-myc-Acn$^{RSB}$] $^{43A1}$; *Sb* / +<br>$w^{*}$ ; GMR-Gal4/ P[w+, uas-myc-Acn$^{SDR}$] $^{43A1}$; *Sb* / + |
| **Figure 3** | **Genotype** |
| 3 A,D,G,P,T,Y,AC | $w^{*}$ ; *acn*$^{27}$; P[w$^{+}$, *acn*P-Myc-Acn$^{WT}$]$^{96F3}$ |
| 3 B,E,H,Q,U,Z,AD | $w^{*}$ ; *acn*$^{27}$; P[w$^{+}$, *acn*P-Myc-Acn$^{NLS}$]$^{96F3}$ |
| 3 C,F,I,R,V,AA,AE | $w^{*}$ ; *acn*$^{27}$; P[w$^{+}$, *acn*P-Myc-Acn$^{NES}$]$^{96F3}$ |
| 3 J | $w^{*}$ ; *acn*$^{27}$; P[w$^{+}$, *acn*P-Myc-Acn$^{WT}$]$^{96F3}$<br>$w^{*}$ ; *acn*$^{27}$; P[w$^{+}$, *acn*P-Myc-Acn$^{NLS}$]$^{96F3}$<br>$w^{*}$ ; *acn*$^{27}$; P[w$^{+}$, *acn*P-Myc-Acn$^{NES}$]$^{96F3}$ |
| 3 K | $w^{*}$ ; GMR-Gal4 /+ |
| 3 L | $w^{*}$ ; GMR-Gal4, P[w+, pAttb-uas-Myr-Acn]/ CyO |
| 3 M | $w^{*}$ ; GMR-Gal4 /+<br>$w^{*}$ ; GMR-Gal4, P[w+, pAttb-uas-Myr-Acn]/ CyO |
| 3 N,S,W,AB,AF | $w^{*}$ ; *acn*$^{27}$/ CyO; P[w$^{+}$, *acn*P-Myc-Acn$^{Myr}$]$^{96F3}$/+ |
| 3 O | $w^{*}$ ; *acn*$^{27}$; P[$w^{+}$, *acn*P-Myc-Acn$^{WT}$]$^{96F3}$<br>$w^{*}$ ; *acn*$^{27}$/ CyO; P[w$^{+}$, *acn*P-Myc-Acn$^{Myr}$]$^{96F3}$/+ |
| 3 X,AG | $w^{*}$ ; *acn*$^{27}$; P[$w^{+}$, *acn*P-Myc-Acn$^{WT}$]$^{96F3}$<br>$w^{*}$ ; *acn*$^{27}$; P[w$^{+}$, *acn*P-Myc-Acn$^{NLS}$]$^{96F3}$<br>$w^{*}$ ; *acn*$^{27}$; P[w$^{+}$, *acn*P-Myc-Acn$^{NES}$]$^{96F3}$<br>$w^{*}$ ; *acn*$^{27}$/ CyO; P[w$^{+}$, *acn*P-Myc-Acn$^{Myr}$]$^{96F3}$/+ |
| 3 AH | $w^{*}$ ; GMR-Gal4 /+ |
| 3 AI | $w^{*}$ ; GMR-Gal4, P[w+, uas-myc-Acn$^{WT}$] / CyO |
| 3 AJ | $w^{*}$ ; GMR-Gal4, P[w+, uas-myc-Acn$^{D527A}$] / CyO |
| 3 AK | $w^{*}$ ; GMR-Gal4, P[w+, uas-Myr-Acn]$^{43A1/}$ CyO |

| **Figure 4** | **Genotype** |
|---|---|
| 4 A,B | w$^{1118}$<br>*w*$^{*}$ ; CarT-HA-T2A-QF2, P[w$^{+}$, Quas-µID]$^{43A1}$<br>*w*$^{*}$ ; CarT-HA-T2A-QF2, P[w$^{+}$, Quas-µID-Acn$^{NES}$]$^{43A1}$<br>*w*$^{*}$ ; CarT-HA-T2A-QF2, P[w$^{+}$, Quas-µID-Acn$^{SDR}$]$^{43A1}$<br>*w*$^{*}$ ; P[w$^{+}$, uas-Myc-Acn$^{WT}$]$^{43A1}$ / +; P[w+, nSyb-GAL4.P]$^{68A}$ / + |
| 4 C | *w*$^{*}$ ; CarT-HA-T2A-QF2, P[w$^{+}$, Quas-µID]$^{43A1}$<br>*w*$^{*}$ ; CarT-HA-T2A-QF2, P[w$^{+}$, Quas-µID-Acn$^{NES}$]$^{43A1}$ |
| 4 D | *w*$^{*}$ ; CarT-HA-T2A-QF2, P[w$^{+}$, Quas-µID]$^{43A1}$<br>*w*$^{*}$ ; CarT-HA-T2A-QF2, P[w$^{+}$, Quas-µID-Acn$^{SDR}$]$^{43A1}$ |
| 4 E | *w*$^{*}$ ; CarT-HA-T2A-QF2, P[w$^{+}$, Quas-µID-Acn$^{NES}$]$^{43A1}$<br>*w*$^{*}$ ; CarT-HA-T2A-QF2, P[w$^{+}$, Quas-µID-Acn$^{SDR}$]$^{43A1}$ |

| **Figure 5** | **Genotype** |
|---|---|
| 5 A | *w*$^{*}$ ; GMR-Gal4, P[w$^{+}$, uas-Myr-Acn]$^{43A1}$/ CyO |
| 5 B | *w*$^{*}$ ; GMR-Gal4, P[w$^{+}$, uas-Myr-Acn]$^{43A1}$ / *Rab 32*-ltd1 |
| 5 C | *w*$^{*}$ ; GMR-Gal4, P[w$^{+}$, uas-Myr-Acn]$^{43A1}$ / +; *Lrrk*$^{ex1}$/ + |
| 5 D | *w*$^{*}$ ; GMR-Gal4, P[w$^{+}$, uas-Myr-Acn]$^{43A1}$ / CyO |
| 5 E | *w*$^{*}$ ; GMR-Gal4, P[w$^{+}$, uas-Myr-Acn]$^{43A1}$ / uas-Rab32 RNAi (BS 38956); +/ + |
| 5 F | *w*$^{*}$ ; GMR-Gal4, P[w$^{+}$, uas-Myr-Acn]$^{43A1}$ / uas-Lrrk RNAi (BS 39019); +/ + |
| 5 G | OreR |
| 5 H | *w*$^{*}$ ; *acn*$^{27}$/ CyO; P[*w*$^{+}$, *acn*P-Myc-Acn$^{Myr}$]$^{96F3}$/ TM6B |
| 5 I | *w*$^{*}$ ; *Rab 32*-ltd1; P[*w*$^{+}$, *acn*P-Myc-Acn$^{Myr}$]$^{96F3}$/TM6B |
| 5 J | *w*$^{*}$ ; GMR-Gal4 /+ |
| 5 K | *w*$^{*}$ ; GMR-Gal4, P[w+, uas-Myr-Acn]$^{43A1}$ / CyO |
| 5 L | *w*$^{*}$ ; GMR-Gal4, P[w+, uas-Myr-Acn]$^{43A1}$ / CyO; *Lrrk*$^{ex1}$ |
| 5 M,Q,O,S | *w*$^{1118}$ |
| 5 N,R,P,T | *Lrrk*$^{ex1}$ (BS 34750) |
| 5 U | *w*$^{*}$ ; GMR-Gal4, P[w$^{+}$, uas-Myr-Acn]$^{43A1}$/ CyO<br>*w*$^{*}$ ; GMR-Gal4, P[w$^{+}$, uas-Myr-Acn]$^{43A1}$ / *Rab 32*-ltd1<br>*w*$^{*}$ ; GMR-Gal4, P[w$^{+}$, uas-Myr-Acn]$^{43A1}$ / +; *Lrrk*$^{ex1}$/ + |
| 5 V | *w*$^{*}$ ; GMR-Gal4, P[w+, uas-Myr-Acn]$^{43A1}$ / CyO; +/ +<br>*w*$^{*}$ ; GMR-Gal4, P[w+, uas-Myr-Acn]$^{43A1}$ / uas-Rab32 RNAi (BS 38956); +/ +<br>*w*$^{*}$ ; GMR-Gal4, P[w+, uas-Myr-Acn]$^{43A1}$ / uas-Lrrk RNAi (BS 39019); +/ + |
| 5 W | OreR<br>*w*$^{*}$ ; *acn*$^{27}$/ CyO; P[*w*$^{+}$, *acn*P-Myc-Acn$^{Myr}$]$^{96F3}$/ TM6B<br>*w*$^{*}$ ; *Rab 32*$^{ltd-1}$; P[*w*$^{+}$, *acn*P-Myc-Acn$^{Myr}$]$^{96F3}$/TM6B |
| 5 X | *w*$^{*}$ ; GMR-Gal4 /+<br>*w*$^{*}$ ; GMR-Gal4, P[w+, uas-Myr-Acn]$^{43A1}$ / CyO<br>*w*$^{*}$ ; GMR-Gal4, P[w+, uas-Myr-Acn]$^{43A1}$ / CyO; *Lrrk*$^{ex1}$ |
| 5 Y | *w*$^{1118}$<br>*Lrrk*$^{ex1}$ |

### Supplemental Figure 2. Sequences

#### A) µID-Acn fusion proteins

**µID-Acn^SDR**

MGFKNLIWLKEVDSTQERLKEWNVSYGTALVADRQTRGRGGSGRRWFSQEGGLYFSFLLNPKEFENLLQLPLVL
GLSVSEALEEITEIPFSLKWPNDVYFQERKVSGVLCELSKDKLIVGIGINVNQREIPEEIKDRATTLYEITGKDWDRKEVLLK
VLKRISENLKKFKEASDYKDDDDKGTLPPLERLTLTGFMRRRSERKKSSSPAPVPVAVARRSRRSRKVSENEKTEVETPVV
AQNNEAECPKVPPASQEINEARASSEERAGSGASPVRRSRGSRTTPRKRTESKGGDKKVDTIPEEEPENGKDVEKDAVE
EEKTVAPPAANESESEDRAKTEDQQAQDDESCAEPAVVKDDDVEKEDLKPQQEVPTSTDDDTKQSEPEQEVKVPSPSS
KDEEHLKTTSDVLDIQTEEVDERDEEASAPAERRETKKRHKEQIKEQQPAEEEEHKENHKQIDENPEQIHDEPADLSNDK
HDPCLNSSATNTTGAPVLQQQRSVRKRKWLTNKKSSDRTPPVLAISTDSLKNIIADVVQPVPLSDVQLESSSEEEGLVTS
DRDSERSASPAPPSEASKDRDNKGNTANTCDTTAEASADNEGAGAAPPAAADTVSAQVSLSKATVSTAAPHIVRD**PDP**
ARNRASHVLYITNLVRPFTVLQLKGLLARTGKIVEEDGFWIDRIKSKCYVAYSTEDEAIETRHALHGVRWPVSNPKCLNV
DFGSRTD**MAR**AILSTKDEAPRYGQENTRDNQQSGNAWSRLEPSDKKPARPVREWDVGKKEPSDHDKHSNDRRRDSK
DRLDSRSRDAERAGQERKRSRDREGRGRDRERERLDRNAHARSRSGSPASKTKKENEPPIRL**ADDLFRKAKGTPCAAA**
**LPA**TPEAIAEKEAFRQKRIEEHKLRIKEREERQKEREKDRDRQRETRRNRSNERRRSRSRERERRRY*

**µID-Acn^NES**

MGFKNLIWLKEVDSTQERLKEWNVSYGTALVADRQTRGRGGSGRRWFSQEGGLYFSFLLNPKEFENLLQLPLVLGLSV
SEALEEITEIPFSLKWPNDVYFQERKVSGVLCELSKDKLIVGIGINVNQREIPEEIKDRATTLYEITGKDWDRKEVLLKVLKRI
SENLKKFKEASDYKDDDDKGTLPPLERLTLTGFMRRRSERKKSSSPAPVPVAVARRSRRSRKVSENEKTEVETPVVAQN
NEAECPKVPPASQEINEARASSEERAGSGASPVRRSRGSRTTPRKRTESKGGDKKVDTIPEEEPENGKDVEKDAVEEEKT
VAPPAANESESEDRAKTEDQQAQDDESCAEPAVVKDDDVEKEDLKPQQEVPTSTDDDTKQSEPEQEVKVPSPSSKDEE
HLKTTSDVLDIQTEEVDERDEEASAPAERRETKKRHKEQIKEQQPAEEEEHKENHKQIDENPEQIHDEPADLSNDKHDP
CLNSSATNTTGAPVLQQQRSVRKRKWLTNKKSSDRTPPVLAISTDSLKNIIADVVQPVPLSDVQLESSSEEEGLVTSDRDS
ERSASPAPPSEASKDRDNKGNTANTCDTTAEASADNEGAGAAPPAAADTVSAQVSLSKATVSTAAPHIVRDPSPARNR
ASHVLYITNLVRPFTVLQLKGLLARTGKIVEEDGFWIDRIKSKCYVAYSTEDEAIETRHALHGVRWPVSNPKCLNVDFGSR
TDMDRAILSTKDEAPRYGQENTRDNQQSGNAWSRLEPSDKKPARPVREWDVGKKEPSDHDKHSNDRRRDSKDRLDS
RSRDAERAGQERKRSRDREGRGRDRERERLDRNAHARSRSGSPASKTKKENEPPIRLLDDLFRKTKGTPCIYWLPLTPEA
IAEKEAFRQKR**AEEHKARA**KEREERQKEREKDRDRQRETRRNRSNERRRSRSRERERRRY*

**Legend:** µID (Johnson et al., 2022); **FLAG** epitope tag,
Acn: **S437D**; **D527A**; **RSB mutation** NES mutant: **AEEHKARA**

### B) Myristoylated Acn proteins

**uas-Myr-Acn**

MGSSKSKPKDPSQRRRDYKDDDDKDYKDDDDKDYKDDDDKEFMRRRSERKKSSSPAPVPVAVARRSRRSRKVSENE
KTEVETPVVAQNNEAECPKVPPASQEINEARASSEERAGSGASPVRRSRGSRTTPRKRTESKGGDKKVDTIPEEEPENGK
DVEKDAVEEEKTVAPPAANESESEDRAKTEDQQAQDDESCAEPAVVKDDDVEKEDLKPQQEVPTSTDDDTKQSEPEQ
EVKVPSPSSKDEEHLKTTSDVLDIQTEEVDERDEEASAPAERRETKKRHKEQIKEQQPAEEEEHKENHKQIDENPEQIHD
EPADLSNDKHDPCLNSSATNTTGAPVLQQQRSVRKRKWLTNKKSSDRTPPVLAISTDSLKNIIADVVQPVPLSDVQLESS
SEEEGLVTSDRDSERSASPAPPSEASKDRDNKGNTANTCDTTAEASADNEGAGAAPPAAADTVSAQVSLSKATVSTAA
PHIVRDPSPARNRASHVLYITNLVRPFTVLQLKGLLARTGKIVEEDGFWIDRIKSKCYVAYSTEDEAIETRHALHGVRWPV
SNPKCLNVDFGSRTDMDRAILSTKDEAPRYGQENTRDNQQSGNAWSRLEPSDKKPARPVREWDVGKKEPSDHDKHS
NDRRRDSKDRLDSRSRDAERAGQERKRSRDREGRGRDRERERLDRNAHARSRSGSPASKTKKENEPPIRLLDDLFRKTK
GTPCIYWLPLTPEAIAEKEAFRQKRIEEHKLRIKEREERQKEREKDRDRQRETRRNRSNERRRSRSRERERRRY*

**genomic Myr-Acn**

MGSSKSKPKDPSQRRRPAGELAAAMRRRSERKKSSSPAPVPVAVARRSRRSRKVSENEKTEVETPVVAQNNEAECPK
VPPASQEINEARASSEERAGSGASPVRRSRGSRTTPRKRTESKGGDKKVDTIPEEEPENGKDVEKDAVEEEKTVAPPAAN
ESESEDRAKTEDQQAQDDESCAEPAVVKDDDVEKEDLKPQQEVPTSTDDDTKQSEPEQEVKVPSPSSKDEEHLKTTSD
VLDIQTEEVDERDEEASAPAERRETKKRHKEQIKEQQPAEEEEHKENHKQIDENPEQIHDEPADLSNDKHDPCLNSSAT
NTTGAPVLQQQRSVRKRKWLTNKKSSDRTPPVLAISTDSLKNIIADVVQPVPLSDVQLESSSEEEGLVTSDRDSERSASP
APPSEASKDRDNKGNTANTCDTTAEASADNEGAGAAPPAAADTVSAQVSLSKATVSTAAPHIVRDPSPARNRASHVLY
ITNLVRPFTVLQLKGLLARTGKIVEEDGFWIDRIKSKCYVAYSTEDEAIETRHALHGVRWPVSNPKCLNVDFGSRTDMDR
AILSTKDEAPRYGQENTRDNQQSGNAWSRLEPSDKKPARPVREWDVGKKEPSDHDKHSNDRRRDSKDRLDSRSRDA
ERAGQERKRSRDREGRGRDRERERLDRNAHARSRSGSPASKTKKENEPPIRLLDDLFRKTKGTPCIYWLPLTPEAIAEKEA
FRQKRIEEHKLRIKEREERQKEREKDRDRQRETRRNRSNERRRSRSRERERRRY*

**Legend:src Myristoylation signal**; **3xFlag epitope**; Acn

#### C) Gal4/uas-regulated Myc-Acn mutants

**uas-Myc-Acn$^{RSB}$**

MEEQKLISEEDLLEEQKLISEEDLLSTAEFMRRRSERKKSSSPAPVPVAVARRSRRSRKVSENEKTEVETPVVAQNNEAEC
PKVPPASQEINEARASSEERAGSGASPVRRSRGSRTTPRKRTESKGGDKKVDTIPEEEPENGKDVEKDAVEEEKTVAPPA
ANESESEDRAKTEDQQAQDDESCAEPAVVKDDDVEKEDLKPQQEVPTSTDDDTKQSEPEQEVKVPSPSSKDEEHLKTT
SDVLDIQTEEVDERDEEASAPAERRETKKRHKEQIKEQQPAEEEEHKENHKQIDENPEQIHDEPADLSNDKHDPCLNSS
ATNTTGAPVLQQQRSVRKRKWLTNKKSSDRTPPVLAISTDSLKNIIADVVQPVPLSDVQLESSSEEEGLVTSDRDSERSAS
PAPPSEASKDRDNKGNTANTCDTTAEASADNEGAGAAPPAAADTVSAQVSLSKATVSTAAPHIVRDPSPARNRASHVL
YITNLVRPFTVLQLKGLLARTGKIVEEDGFWIDRIKSKCYVAYSTEDEAIETRHALHGVRWPVSNPKCLNVDFGSRTDMD
RAILSTKDEAPRYGQENTRDNQQSGNAWSRLEPSDKKPARPVREWDVGKKEPSDHDKHSNDRRRDSKDRLDSRSRD
AERAGQERKRSRDREGRGRDRERERLDRNAHARSRSGSPASKTKKENEPPIRL**ADDLFRKAKGTPCAAALPA**TPEAIAE
KEAFRQKRIEEHKLRIKEREERQKEREKDRDRQRETRRNRSNERRRSRSRERERRRY*

**uas-Myc-Acn$^{SDR}$**

MEEQKLISEEDLLEEQKLISEEDLLSTAEFMRRRSERKKSSSPAPVPVAVARRSRRSRKVSENEKTEVETPVVAQNNEAEC
PKVPPASQEINEARASSEERAGSGASPVRRSRGSRTTPRKRTESKGGDKKVDTIPEEEPENGKDVEKDAVEEEKTVAPPA
ANESESEDRAKTEDQQAQDDESCAEPAVVKDDDVEKEDLKPQQEVPTSTDDDTKQSEPEQEVKVPSPSSKDEEHLKTT
SDVLDIQTEEVDERDEEASAPAERRETKKRHKEQIKEQQPAEEEEHKENHKQIDENPEQIHDEPADLSNDKHDPCLNSS
ATNTTGAPVLQQQRSVRKRKWLTNKKSSDRTPPVLAISTDSLKNIIADVVQPVPLSDVQLESSSEEEGLVTSDRDSERSAS
PAPPSEASKDRDNKGNTANTCDTTAEASADNEGAGAAPPAAADTVSAQVSLSKATVSTAAPHIVRD**PDP**ARNRASHV
LYITNLVRPFTVLQLKGLLARTGKIVEEDGFWIDRIKSKCYVAYSTEDEAIETRHALHGVRWPVSNPKCLNVDFGSRTD**M**
**AR**AILSTKDEAPRYGQENTRDNQQSGNAWSRLEPSDKKPARPVREWDVGKKEPSDHDKHSNDRRRDSKDRLDSRSR
DAERAGQERKRSRDREGRGRDRERERLDRNAHARSRSGSPASKTKKENEPPIRL**ADDLFRKAKGTPCAAALPA**TPEAIA
EKEAFRQKRIEEHKLRIKEREERQKEREKDRDRQRETRRNRSNERRRSRSRERERRRY*

**Legend: 2xMyc epitope; Acn; S437D; D527A; RSB mutation**

### D) Acinus mutants under control of Acn promoter

**gen-Myc-Acn$^{NLS}$**

MEEQKLISEEDLLEEQKLISEEDLLSTAEFMRRRSERKKSSSPAPVPVAVARRSRRSRKVSENEKTEVETPVVAQNNEAEC
PKVPPASQEINEARASSEERAGSGASPVRRSRGSRTTPRKRTESKGGDKKVDTIPEEEPENGKDVEKDAVEEEKTVAPPA
ANESESEDRAKTEDQQAQDDESCAEPAVVKDDDVEKEDLKPQQEVPTSTDDDTKQSEPEQEVKVPSPSSKDEEHLKTT
SDVLDIQTEEVDERDEEASAPAERRETKKRHKEQIKEQQPAEEEEHKENHKQIDENPEQIHDEPADLSNDKHDPCLNSS
ATNTTGAPVLQQQRSVRKRKWLTNKKSSDRTPPVLAISTDSLKNIIADVVQPVPLSDVQLESSSEEEGLVTSDRDSERSAS
PAPPSEASKDRDNKGNTANTCDTTAEASADNEGAGAAPPAAADTVSAQVSLSKATVSTAAPHIVRDPSPARNRASHVL
YITNLVRPFTVLQLKGLLARTGKIVEEDGFWIDRIKSKCYVAYSTEDEAIETRHALHGVRWPVSNPKCLNVDFGSRTDMD
RAILSTKDEAPRYGQENTRDNQQSGNAWSRLEPSDKKPARPVREWDVGKKEPSDHDKHSNDRRRDSKDRLDSRSRD
AERAGQERKRSRDREGRGRDRERERLDRNAHARSRSGSPASKTKKENEPPIRLLDDLFRKTGTPCIYWLPLTPEAIAEKE
AFRQKRIEEHKLRIKEREERQKEREKDRDRQRETRRNRSNEAAASASAEAEARRY*

**gen-Myc-Acn$^{NES}$**

MEEQKLISEEDLLEEQKLISEEDLLSTAEFMRRRSERKKSSSPAPVPVAVARRSRRSRKVSENEKTEVETPVVAQNNEAEC
PKVPPASQEINEARASSEERAGSGASPVRRSRGSRTTPRKRTESKGGDKKVDTIPEEEPENGKDVEKDAVEEEKTVAPPA
ANESESEDRAKTEDQQAQDDESCAEPAVVKDDDVEKEDLKPQQEVPTSTDDDTKQSEPEQEVKVPSPSSKDEEHLKTT
SDVLDIQTEEVDERDEEASAPAERRETKKRHKEQIKEQQPAEEEEHKENHKQIDENPEQIHDEPADLSNDKHDPCLNSS
ATNTTGAPVLQQQRSVRKRKWLTNKKSSDRTPPVLAISTDSLKNIIADVVQPVPLSDVQLESSSEEEGLVTSDRDSERSAS
PAPPSEASKDRDNKGNTANTCDTTAEASADNEGAGAAPPAAADTVSAQVSLSKATVSTAAPHIVRDPSPARNRASHVL
YITNLVRPFTVLQLKGLLARTGKIVEEDGFWIDRIKSKCYVAYSTEDEAIETRHALHGVRWPVSNPKCLNVDFGSRTDMD
RAILSTKDEAPRYGQENTRDNQQSGNAWSRLEPSDKKPARPVREWDVGKKEPSDHDKHSNDRRRDSKDRLDSRSRD
AERAGQERKRSRDREGRGRDRERERLDRNAHARSRSGSPASKTKKENEPPIRLLDDLFRKTGTPCIYWLPLTPEAIAEKE
AFRQKR**AEEHKARA**KEREERQKEREKDRDRQRETRRNRSNERRRSRSRERERRRY*

**Legend:**

**2xMyc epitope; NLS mutation AAASASAEAEA; NES mutation AEEHKARA**

### E) PCR primers

#### Piwi RT-PCR

Piwi_ex4fwd: AATTCCTGAGCTCTGCCGAGTG

Piwi_ex5rev: TAACTGCTCATGGCACGCATAA

#### ***Rab32***$^{ltd-1}$ **genotyping** (PCR followed by sequencing)

Rab32_fwd: AAGCGCCCAGACTATTGCTA

Rab32_rev: GGTCATGTTGCCAAACCTCT

#### ***Lrrk***$^{ex1}$ **genotyping**

Lrrk_fwd: ATGGAGTTTCCGGACATCAG

Lrrk_rev: CGTTTGCATTTTCGTGCTT
